# Brain-wide circuitry underlying altered auditory habituation in zebrafish models of autism

**DOI:** 10.1101/2024.09.04.611137

**Authors:** Maya Wilde, Anahita Ghanbari, Tessa Mancienne, Ailís Moran, Rebecca E. Poulsen, Lena Constantin, Conrad Lee, Leandro Aluisio Scholz, Joshua Arnold, Wei Qin, Timothy J. Karle, Steven Petrou, Itia Favre-Bulle, Ellen J. Hoffman, Ethan K. Scott

## Abstract

Auditory processing is widely understood to occur differently in autism, though the patterns of brain activity underlying these differences are not well understood. The diversity of autism also means brain-wide networks may change in various ways to produce similar behavioral outputs. We used larval zebrafish to investigate auditory habituation in four genetic lines relevant to autism: *fmr1*, *mecp2*, *scn1lab* and *cntnap2*. In free-swimming behavioral tests, we found each line had a unique profile of auditory hypersensitivity and/or delayed habituation. Combining the optical transparency of larval zebrafish with genetically encoded calcium indicators and light-sheet microscopy, we then observed brain-wide activity at cellular resolution during auditory habituation. As with behavior, each line showed unique alterations in brain-wide spontaneous activity, auditory processing, and adaptation in response to repetitive acoustic stimuli. We also observed commonalities in activity across our genetic lines that indicate shared circuit changes underlying certain aspects of their behavioral phenotypes. These were predominantly in regions involved in sensory integration and sensorimotor gating rather than primary auditory areas. Overlapping phenotypes include differences in the activity and functional connectivity of the telencephalon, thalamus, dopaminergic regions, and the locus coeruleus, and excitatory/inhibitory imbalance in the cerebellum. Unique phenotypes include loss of activity in the habenula in *scn1lab*, increased activity in auditory regions in *fmr1,* and differences in network activity over time in *mecp2* and *cntnap2*. Comparing these distinct but overlapping brain-wide auditory networks furthers our understanding of how diverse genetic factors can produce similar behavioral effects through a range of circuit- and network-scale mechanisms.

## Introduction

### Sensory processing in autism

Differences in sensory experience are common in autism, but the differences in neural circuitry underlying these traits is not well understood^1–5^. Human studies provide conflicting results about various measures of auditory processing in autism, and these discrepancies may arise from the etiological complexity of autism ^6–8^. More consistent results can be found in studies of syndromic forms of autism, such as fragile X syndrome (FXS) or Rett syndrome, but this approach risks poor representation of autism as a whole ^7,9,10^. There is evidence for reduced habituation to auditory stimuli in some autistic people and reduced adaptation of auditory cortex activity in FXS ^8,11^. There are several hypotheses explaining differences in brain activity in autism, including excitatory/inhibitory (E/I) imbalance, dopaminergic dysfunction, and altered cerebellar and brainstem function ^12–15^. It is not clear which of these link to changes in auditory habituation, or indeed whether different mechanisms are relevant to different etiologies. Animal models enable more incisive studies into brain function, but auditory habituation is under-studied in rodent models of autism compared to other measures of auditory function, despite its simplicity and potential to affect conclusions about differences in other auditory tests ^8,16,17^. While most animal models of autism only manipulate single genes, comparing across models can provide insights into the diversity of mechanisms underlying shared behavioral changes ^16,18^.

### Zebrafish genetic lines for investigating brain-wide function

There is growing interest in using zebrafish to investigate differences in brain development in autism due to their genetic tractability and capacity for brain-wide calcium imaging ^19–21^. They enable investigations of responses to a range of different sensory stimuli including visual, acoustic, vestibular, olfactory, and water flow^22–30^. Indeed, differences in sensory processing in *fmr1^-/-^* fish have illustrated the advantages of capturing cellular resolution activity throughout the brain to describe phenotypes, with phenotypes characterized by differences in functional connectivity rather than in gross activity level ^31,32^. Furthermore, the relative efficiency of zebrafish research supports the shift in autism research towards comparing phenotypes across several different animal lines to better capture the complexity of autism etiology^16,17,21,33,34^.

Auditory habituation is well established in larval zebrafish, and pharmacological and optogenetic manipulations have found roles for dopamine, serotonin, glycine, and NMDA receptors in this process ^35–40^. Increased dopamine signaling increases the degree of habituation, while decreased dopamine signaling reduces the rate and degree of habituation ^38,39^. Serotonin has the opposite role to dopamine: increased serotonin reduces the degree of habituation while reducing serotonergic activity, particularly from the superior raphe, increases habituation^38^. In one study, glycine receptor blocker strychnine entirely eliminated habituation^39^. Blocking NMDA receptors also reduces the rate and/or degree of habituation^35,36,39,40^. Of note, reduced NMDAR activity has been linked to auditory phenotypes in rodent models of autism^41,42^.

### Distinct but overlapping mechanisms for altered habituation

The reduction in response during habituation is generally viewed as a learned association to the innocuous nature of a stimulus. However, a superficially similar phenomenon of ‘induced passivity’ represents a different process: learning that behavioral responses are futile for escaping the stimulus. The networks underlying these two processes may or may not overlap, but both can be disrupted with NMDAR antagonist ketamine ^36,43^. Interpreting this induced passivity as behavioral adaptation fits with the newer narrative that habituation involves shifting response strategy, and is more complex than simply ‘learning to ignore’ stimuli ^44^. A putative passivity circuit in larval zebrafish has been described in the context of electric shock stimuli, where passivity was linked to increased activity in the ventral habenula and decreased activity in the dorsal thalamus and superior raphe^43^. These neurotransmitters and brain regions present diverse potential mechanisms for auditory habituation phenotypes in neurological conditions such as autism.

Here, we set out to study auditory habituation in four genetic lines associated with autism: *fmr1*, *scn1lab, mecp2*, and *cntnap2*, using both behavioral screening and imaging of whole-brain activity at single-neuron resolution. Each of these genes is associated with autism as well as its own syndrome: *fmr1* with FXS, *mecp2* with Rett syndrome, *scn1lab* with Dravet syndrome, and *cntnap2* with Pitt-Hopkins like syndrome^45^. We find that each line has a unique behavioral phenotype in response to repetitive acoustic stimuli, and that they show distinct but overlapping changes in their brain-wide auditory networks during habituation. The results give a glimpse of the behavioral and functional complexity of autism-associated genes and identify network-scale alterations that could contribute to sensory changes in specific syndromic forms of autism.

## Methods

### Animals

All experiments were conducted on 6 days post fertilization zebrafish larvae on a Tüpfel-Longfin background. For imaging of fluorescent calcium transients, *mitfa^-/-^* fish transgenic for *HuC:H2B- GCaMP6s* were used^46^. Larvae were raised in embryo media (distilled water with 10% Hanks solution, consisting of 137mM NaCl, 5.4mM KCl, 0.25mM Na2HPO4, 0.44mM KH2PO4, 1.3mM CaCl2, 1.0mM 654 MgSO4 and 4.2mM NaHCO3 at pH 7.2) in an incubator at 28°C with a 14/10 hour light/dark cycle. The experimental room was maintained at approximately 26°C. Experimental animals were bred from parents heterozygous for the relevant autism-associated gene mutation to provide sibling wild-type controls for each dataset. Separate datasets were collected for each of the four lines: *fmr1^hu27^*^87^, *mecp2^fh2^*^32^, *scn1lab^Δ^*^44^ ^21,47^, and *cntnap2a^ya2188^cntnap2b^ya20^*^43^ ^48^. As *cntnap2* is duplicated in zebrafish, parents of the experimental animals for the *cntnap2* dataset were heterozygotes for both *cntnap2a* and *cntnap2b*. Following experiments, larvae were euthanized with ice and digested in 100 µL TE buffer with 1 µL ProK (New England Biolabs). They were then genotyped by PCR and Sanger sequencing. Primer sequences for each PCR can be found in Supplementary table 1. Larvae were genotyped after the experiment to maintain experimental blindness.

### Free-swimming auditory habituation

#### Experimental set-up

Initial behavioral phenotyping was conducted as part of a larger sensory phenotype screening procedure which also involved visual stimuli (data not shown). Free-swimming behavioral experiments were conducted on a custom-built behavioral rig (Figure 1A). Due to lack of swim bladder inflation in *scn1lab^-/-^* fish, for this mutant line all fish were partially embedded upright in 2% low melting point agarose, and behavioral responses assessed based on tail deflections. Fish were placed in seven individual circular wells of diameter 20 mm, arranged around a central speaker glued to the underside of the well plate. The speaker was driven by an amplifier (Dayton Audio DA30 2 × 15W Class D Bridgeable Mini Amplifier), which received input directly from the MATLAB code driving the experiment. The wells were illuminated from below by an array of infrared LEDs (840 nm). A projector delivered visual stimuli to an angled cold mirror to provide constant medium grey background light (500 lx) to the fish, and to deliver visual stimuli. Videos were recorded with a high-speed camera (Ximea xiB-64 model CB019MG-LX-X8G3), with an infrared filter, and with an exposure time of 1 ms and a framerate of 100 fps.

**Figure 1:**
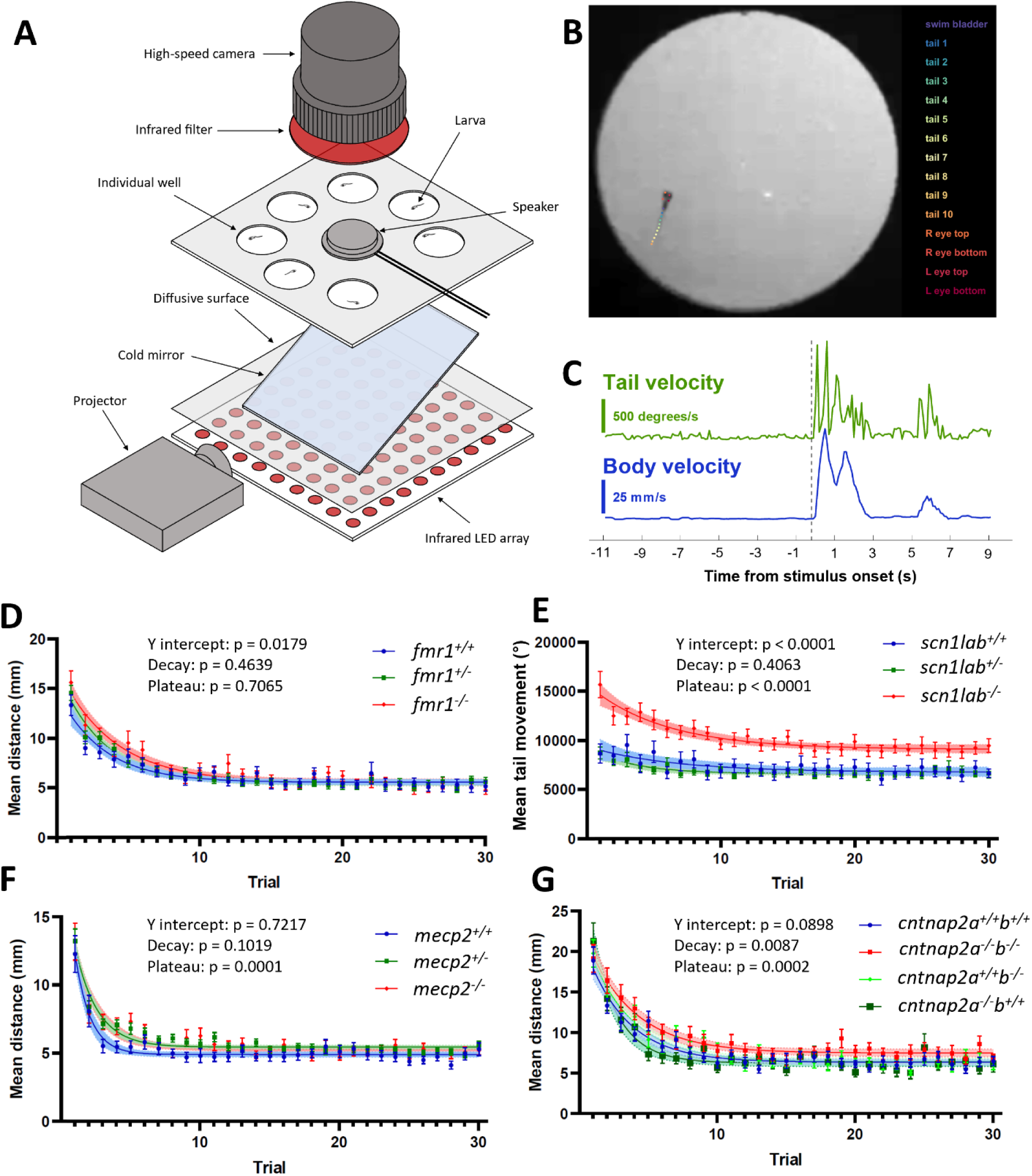
Behavioral auditory habituation phenotypes in four genetic lines. A) Experimental set-up for recording behavior of seven larvae simultaneously. B) Tracking with DeepLabCut^43^. Automated identification of points in the swim bladder, eyes, and along the tail enables kinematic analysis of behavioral responses. C) Example outputs of tail velocity (green) and whole-body velocity (blue) in response to an auditory stimulus, indicated by the dotted line. D-G) Behavioral responses during habituation for *fmr1*, *scn1lab*, *mecp2*, and *cntnap2*, respectively. Responses are calculated for a 1- second window after the stimulus. For each group, the p-value is indicated for y-intercept (Y0), slope, and plateau, comparing fish carrying the mutation to wild-type siblings.

#### Stimuli

The auditory habituation stimulus train was preceded by stimuli for screening of other audiovisual phenotypes, including sounds at different volumes and looming stimuli. Sound stimuli for the auditory habituation test were 500 ms white noise bursts at 108 dBSPL, with 2 ms on and off ramps. The interstimulus interval was 5 seconds, and stimuli were divided into two blocks. The first block consisted of 30 stimuli, followed by a 90 second break, then the recovery block consisted of 15 stimuli.

#### Analysis

Videos were segmented using python (version 3.7.6), and the activity of fish was tracked using DeepLabCut^49^, with a model trained on in-house data for larval zebrafish (Figure 1B-C). Responses were analyzed with custom MATLAB (2022) scripts. In the *scn1lab* dataset where the fish were partially immobilized, only the tail activity was analyzed. Responses were summed within a 1-second window after the stimulus onset.

Response intensities were compared between genotypes with a non-linear regression to a one-phase decay in GraphPad Prism (v9.4.1). This enabled comparison of whether each dataset can be explained by curves with the same coefficients in the equation 𝑦𝑦 = 𝑎𝑎𝑒𝑒^𝑏𝑏𝑏𝑏^ + 𝑐𝑐. These values represent the y-intercept (a, initial response intensity), slope (b, rate of habituation), and plateau (c, extent of habituation).

### Calcium imaging of brain activity during auditory habituation

#### Data acquisition

Imaging of calcium fluorescence was conducted with a custom-built light-sheet microscope ^31,50,51^. Fish were held in a 3D-printed plastic chamber with glass coverslip walls, filled with embryo media (Supplementary Figure 1A). Fish were fully immobilized in 2% low melting point agarose, except for the *mecp2* dataset, where the tail was cut free to allow a range of motion of 90 degrees to each side. This enabled imaging of tail activity using a camera below the experimental chamber, for confirmation that movements otherwise interpreted from motion correction of brain imaging indeed correlated to real tail movements (data not shown).

The brain was imaged by scanning two perpendicular sheets of light through 50 z-planes at a step size of 5 µm, to cover a total volume of 250 µm with a frame rate 100 fps and binning of 4, resulting in volumetric acquisition of 2 Hz (Supplementary Figure 1B). The imaging column was as previously described^50^, consisting of a 20x water immersion objective, a filter to exclude the 488 nm wavelength light from the excitation laser, an electrically tunable lens, and a high-speed camera (PCO edge 5.5). Acquisition and delivery of acoustic stimuli were controlled with Micro Manager software (version 1.4) ^52^, and a custom written GUI in MATLAB (2022).

#### Stimuli

Acoustic stimuli were delivered via a speaker (Dayton Audio DAEX-9-4SM Skinny Mini Exciter Audio, Haptic Item Number 295-256) affixed to the back wall of the experimental chamber, so that sounds were delivered directly into the embryo media filling the chamber^27^. The stimulus train was again part of a wider screening protocol: firstly, a separate recording of 10 minutes of spontaneous activity (data not shown), then a second recording of sounds at different volumes (data not shown), followed by the auditory habituation paradigm. The auditory habituation stimulus train was composed of 100 ms white noise bursts with 2 ms on and off ramps, set to a volume equivalent to 96 dBSPL in air. Due to the requirement for smaller speakers to fit on the experimental chamber, it was not possible to deliver sounds precisely matching those in the free-swimming set-up. The interstimulus interval was 3 seconds, and again the stimuli were broken into two blocks, except for the *scn1lab* dataset, which did not have a recovery block. The first block consisted of 20 stimuli, followed by 10 stimuli in the second block after a 1-minute break.

#### Analysis of neuronal traces

Regions of interest (ROIs) representing individual neurons, and their fluorescent traces over time were extracted using Suite2p (Supplementary Figure 1B)^53^. The mean stack of images was then warped, first to a template brain averaged from 10 wild-type larvae, then to the Zbrain reference brain space, using the ANTs warping algorithm^54,55^. A mask of all the brain regions in this reference atlas was then used to exclude any extraneous ROIs identified outside the brain or in the eyes. The extraction and warping steps were performed using the high-performance computing cluster at the University of Melbourne. The remaining analysis was conducted in MATLAB (version 2022). The ΔF/F of the fluorescent traces was calculated using a sliding window of 201 timepoints, and a smoothing kernel of 7 timepoints.

Correlation to auditory stimuli and motion were calculated by linear regression to theoretical calcium transients at stimulus timings and timings of motion correction as outputted by Suite2p (Supplementary Figure 1C-D). Comparisons of metrics without data for each neuron were performed with Wilcoxon ranked-sum tests. For voxel-wise spatial comparison of various measures, neurons were averaged in 3-dimensional cubes of edge length 10µm. Mean activity traces of neurons within these cubes were fitted to a curve described by the function 𝑦𝑦 = 𝑎𝑎𝑒𝑒^𝑏𝑏𝑏𝑏^ + 𝑐𝑐 to obtain three curve fit parameters.

We performed graph theory analysis with the brain connectivity toolbox for MATLAB^56^. For each fish, a correlation matrix across all neurons was produced based on correlations in ΔF/F activity within the period of interest, and autocorrelations were removed. The matrix was binarized using one of two methods: the top 10% highest correlations, or correlation coefficients above a threshold of 0.3. The degree of each neuron was then calculated as the proportion of the maximum possible number of edges.

We compared a range of metrics between genotypes within anatomical regions, using linear mixed effects models. The equation used was ’Y ∼ genotype + (1|genotype:fishID)’, where Y is a vector of some response metric for each cell within a region. The fixed effect is the genotype, and the random effect is the individual fish ^57^. The region list came from the Zbrain reference atlas, with custom added masks for the octavolateralis nucleus and the granule cells of the cerebellum. We set inclusion criteria for brain regions that at least 60% of wild-type fish from each of the 4 datasets must have at least 5 neurons identified in that region. We further excluded regions in which we would not expect to find cell nuclei, and small regions defined by expression of certain markers. A full list of p-values from all comparisons can be found in the Supplementary Information.

We calculated the ratio between the degree in the *gad1b* and *vglut2* parts of the cerebellum for the *fmr1* and *scn1lab* datasets, excluding any ROIs which overlapped between the two, using the average degree per fish for each time period and a repeated measures ANOVA, with a Dunn-Sidak test for multiple comparisons.

Total motor activity, genotype, and correlation threshold for the top 10% of edges in the *cntnap2* dataset were compared with a linear mixed-effects model with the equation ’Y ∼ motor + (1|genotype)’, where Y is the correlation threshold, the fixed effect is motor activity, and the random effect is the genotype. Each data point is one fish in this analysis.

## Results

### Free-swimming auditory habituation phenotypes

As expected, the behavioral responses to auditory stimuli were well modelled by an exponential decay curve. Within each dataset, we compared fit metrics of the y-intercept (the calculated y value of the curve at the initial stimulus), decay rate (slope), and final plateau value. For the *fmr1* dataset, the y-intercept was higher for the *fmr1^-/-^* (14.50, n = 38) and *fmr1^+/-^* (13.79, n = 77) fish than the *fmr1^+/+^* controls (12.27, n = 41, p = 0.0179, Figure 1D). However, the slope was not different between genotypes (*fmr1^+/+^* 0.319, *fmr1^+/-^* 0.320, *fmr1^-/-^* 0.263, p = 0.4639), and neither was the plateau (*fmr1^+/+^* 5.57, *fmr1^+/-^* 5.43, *fmr1^-/-^* 5.55, p = 0.7065). The *fmr1* mutation therefore produces an initial sensitivity phenotype, but the rate of habituation is the same as wild types, and habituation eventually reaches the same plateau as in wild-type siblings. Therefore, if the degree of habituation is measured as relative to the initial response, the habituation strength could be considered increased in the *fmr1^-/-^* fish.

As *scn1lab^-/-^* larvae do not consistently swim upright, we embedded their heads in agarose and measured their responses based on tail movements rather than distance travelled (Figure 1E). Very striking differences in the mutants were apparent in the y-intercept (*scn1lab^+/+^* 9002, n = 17, *scn1lab^+/-^* 8674, n =47, *scn1lab^-/-^* 14639, n = 31, p < 0.0001) and the plateau (*scn1lab^+/+^* 6769, *scn1lab^+/-^* 6639, *scn1lab^-/-^* 9092, p < 0.0001), but not in the decay rate of the curve (*scn1lab^+/+^* 0.158, *scn1lab^+/-^* 0.317, *scn1lab^-/-^* 0.177, p = 0.4063). The *scn1lab* homozygous mutants therefore have a very strong hypersensitivity phenotype, with a reduced extent of habituation in absolute measures.

In the *mecp2* line (Figure 1F), the y-intercept was not different between genotypes (*mecp2^+/+^* 12.28, n = 40, *mecp2^+/-^* 12.76, n = 107, *mecp2^-/-^* 12.47, n = 50, p = 0.7217), nor was the decay rate (*mecp2^+/+^* 0.907, *mecp2^+/-^* 0.618, *mecp2^-/-^* 0.5714, p = 0.4639). However, the plateau was significantly higher in the *mecp2^-/-^* fish (5.334) and *mecp2^+/-^* fish (5.442) than the *mecp2^+/+^* fish (4.900, p = 0.0001). The *mecp2* phenotype is therefore reduced extent of habituation.

For the *cntnap2* line, both the decay rate (*cntnap2a^+/+^b^+/+^* 0.337, n = 31, *cntnap2a^+/+^b^-/-^* 0.320, n = 25, *cntnap2a^-/-^b^+/+^* 0.519, n = 21, *cntnap2a^-/-^b^-/-^* 0.293, n = 30, p = 0.0087) and the plateau level (*cntnap2a^+/+^b^+/+^* 6.355, *cntnap2a^+/+^b^-/-^* 6.443, *cntnap2a^-/-^b^+/+^* 6.256, *cntnap2a^-/-^b^-/-^* 7.480, p = 0.0002) were significantly different between genotypes (Figure 1G). For the plateau, this difference is driven by the *cntnap2a^-/-^b^-/-^* fish having a higher plateau response rate than the other genotypes. The *cntnap2a^-/-^ b^-/-^* fish have a slower decay rate than the wild types and *cntnap2a^+/+^b^-/-^* fish, but *cntnap2a^-/-^b^+/+^* fish diverge in the other direction, with a faster habituation rate. The y-intercept did not reach the significance threshold (*cntnap2a^+/+^b^+/+^* 17.88, *cntnap2a^+/+^b^-/-^* 18.89, *cntnap2a^-/-^b^+/+^* 21.00, *cntnap2a^-/-^b^-/-^* 19.37, p = 0.0898), but the strongest difference is between the wild type initial response, and the highest initial response of the *cntnap2a^-/-^b^+/+^* fish. Overall, the homozygous mutant of both paralogs habituates both more slowly and to a lesser extent than wild types.

### Brain-wide imaging of auditory habituation phenotypes

#### Brain-wide auditory phenotypes at different scales

##### Whole-brain measures

To understand the brain activity underlying the behavioral auditory habituation phenotypes for each of the four genetic lines, we performed calcium imaging using light-sheet microscopy and the genetically encoded calcium indicator GCaMP6s, expressed in the nuclei of all neurons. This enabled detection of activity at cellular resolution across the full volume of the brain. To quantify activity across the brain-wide network, we used Suite2p^53^ to identify regions of interest (ROIs) generally corresponding to individual neurons^31,58^, and then extracted fluorescence across the experiment for each ROI. The number of ROIs segmented across the whole brain was not different between mutant and wild-type larvae for any of our genetic lines (Supplementary Figure 1E). We next measured correlation to motor activity using the motion correction output from Suite2p (Supplementary Figure 1D). Most movements detected were strong and likely stimulus-evoked rather than spontaneous, as expected in restrained fish without visual feedback ^59^. The amount of motor activity was not different to wild types in *fmr1^-/-^*, *mecp2^-/-^* or *cntap2a^-/-^b^-/-^* fish, but was significantly higher in *scn1lab^-/-^* fish, recapitulating the strong behavioral hypersensitivity (Supplementary Figure 1F). We also measured correlation to auditory stimuli, either only to the auditory habituation train or to all sounds (Supplementary Figure 1C, G-H), since the inclusion of quieter sounds aids in distinguishing auditory-specific from stimulus-evoked motor responses. There were no significant brain-wide differences in auditory correlation in any genetic line, indicating that phenotypes arise at the level of sub-regions within the brain.

To explore whether such small subregions had altered activity in our mutants, we used a voxel-wise subsampling approach, creating 10μm cubes throughout the brain and looking for differences across genotypes for a range of metrics (Figures 2-5, B-C). For each of our genetic lines, this approach revealed a unique profile of regions where activity in mutants diverged from wild types (detailed below).

**Figure 2:**
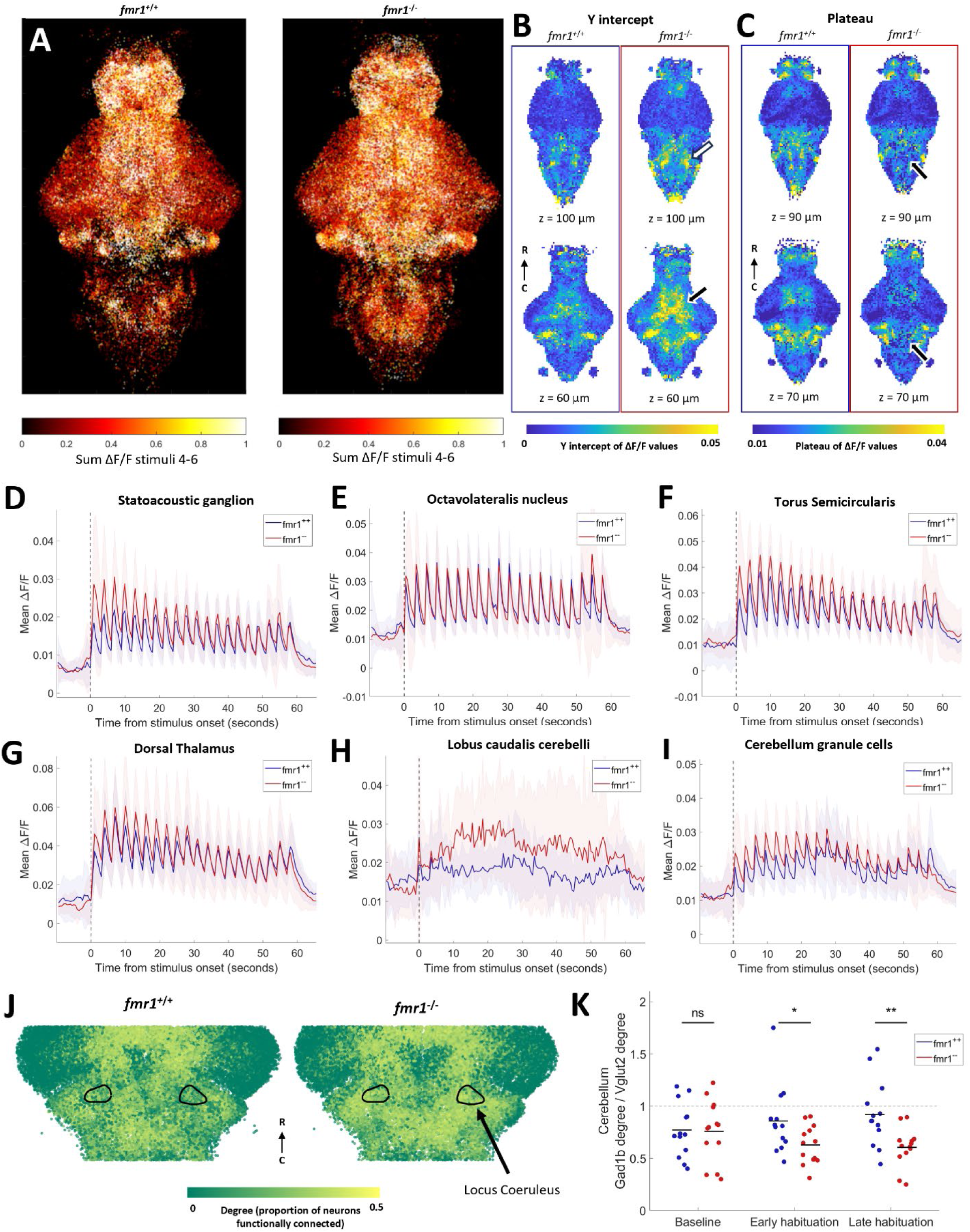
Auditory habituation phenotypes in *fmr1*. A) All segmented neurons from all fish, colored by sum of activity between stimuli 4 and 6. B-C) Comparison within 10 µm cubes of curve fit values during habituation period. Two different z-depths for each measure are shown for *fmr1^+/+^* (left, n = 12) and *fmr1^-/-^* (right, n = 13). B) Y-intercept values are higher in the *fmr1^-/-^* fish, notably in the thalamus (white arrow) and hindbrain (black arrow). C) Plateau values are lower in *fmr1^-/-^* fish than wild types in several parts of the hindbrain (black arrows). D-I) Mean activity of all neurons in the SAG (D), ON (E), TS (F) dorsal thalamus (G) lobus caudalis cerebelli (H), and the granule cell region of the cerebellum (I). D-I: Shading indicates SD. J) A subset of ROIs, colored by degree, identified using the top 10% of edges from correlation during the whole habituation period. Black outlines indicate the locus coeruleus. K) Ratio between the degree of all neurons in *gad1b* and *vglut2* regions of the cerebellum at different periods during habituation. Degree is based on the top 10% of edges. Each dot represents one fish, and black lines indicate the means. Significant effect of genotype (p = 0.0289) and interaction between genotype and time (p = 0.0401), but no significant effect of time alone (p = 0.9167, repeated measures ANOVA). No difference between genotypes at baseline (p = 0.9063), but significantly lower *gad1b*:*vglut2* ratio in *fmr1^-/-^* in the early (p = 0.0380) and late (p = 0.0050) habituation periods (Dunn- Sidak test).

#### Cellular-resolution analyses within anatomical regions

Similarly, we performed analyses within specific brain regions, as defined in the Zbrain atlas^55^. Within these sub-regions we compared several different metrics using cellular-level data, but with a linear mixed effects model that allow us to control for fish of origin^57^. The significance of the effect of genotype across brain regions and metrics are presented in Supplementary Figure 2. These metrics include the auditory and motion correlation as described above, and the three parameters from exponential curve fits to auditory responses. We also compared the number of neurons segmented, and the sum of ΔF/F values in different time periods during the experiment. Lastly, we incorporated the degree measure of functional connectivity, calculated using correlations between all neurons, allowing us to detect differences in network dynamics that would only be evident with cellular resolution. These analyses were intended to characterize changes in brain activity that may contribute to auditory habituation phenotypes.

The summary grids in Supplementary Figure 2 illustrate the distribution of phenotypes throughout the brains of each genotype, with full p-values reported in the Supplementary Material. The *scn1lab^-/-^* fish clearly have many more differences across metrics and brain regions, which is unsurprising given their dramatic behavioral phenotype. None of the phenotypes were common across three or all four genotypes, but there were several instances in which two mutants showed similar effects (Supplementary Figure 2).

#### Brain wide phenotypes in four genetic autism lines

##### Auditory structures are hyperresponsive in *fmr1*

The behavioral phenotype in *fmr1^-/-^* animals was specifically in the y-intercept, representing higher initial responses to acoustic stimuli (Figure 1). Consistent with the behavioral phenotype, there is broadly stronger activity in individual neurons across the brains of *fmr1^-/-^* animals (Figure 2A). When these data are represented using the voxel-based approach, the y-intercept of neuronal activity is also higher in several brain regions, including broad regions across the diencephalon and mesencephalon (Figure 2B ,Supplementary Figure 2A). There is also a portion of the hindbrain that has a lower plateau value compared to wild types (Figure 2C), consistent with the proportionally deeper behavioral habituation observed in mutants.

When single-neuron data are partitioned into defined brain regions, elevated responses are seen throughout the core auditory pathway early in the stimulus train (Figure 2D-G). In the statoacoustic ganglion (SAG), homologous to the cochlear and vestibular ganglia in mammals ^60–62^, the initial response strength is elevated (p = 0.0232 ,Figure 2D), closely mirroring the behavioral hypersensitivity (Figure 1D), before dropping to wild type levels later in the stimulus train. In the octavolateralis nucleus (ON), homologous to the cochlear nucleus^60^ , this effect is not seen across all neurons (p = 0.3454, Figure 2E), but upon closer inspection, strongly responding auditory neurons in the ON have elevated responses early in the stimulus train (Supplementary Figure 3). These results suggest that, while these responses are masked by the large and diverse population of neurons in the ON, there are elevated auditory signals in this structure. Similar elevations in initial response strength are present in the torus semicircularis (TS, p = 0.0352 , Figure 2F), homologous to the inferior colliculus^63^, and the dorsal thalamus (p = 0.0459, Figure 2G), the auditory region within the thalamus^62^.

Beyond the defined auditory processing pathway, cellular-resolution data reveal stronger responses in *fmr1^-/-^* animals for the lobus caudalis (the vestibular region^64^) of the cerebellum (Figure 2H) and a region corresponding to cerebellar granule cells (Figure 2I). The increase in the lobus caudalis does not attenuate like the auditory pathway (Figure 2D-G), showing some degree of elevation throughout the stimulus train. Other observations of cellular-resolution data include increased activity at baseline in the telencephalon of *fmr1^-/-^* fish (p = 0.0434), and an increased number of neurons detected in the pineal (p = 0.0070, Supplementary Figure 2A).

In applying graph theory to data from individual neurons, we found higher functional connectivity (as measured by degree) in the locus coeruleus of *fmr1^-/-^* fish throughout the experiment (Figure 2J, p = 0.0002). In the cerebellum, there are divergent degree differences in the *vglut2*- versus the *gad1b*- enriched areas as habituation proceeds (Supplementary Figure 2). This divergence is of particular interest given their opposing effects on cerebellar output^64,65^. We therefore compared the mean degree of cells within these regions as a ratio over the course of the habituation period (Figure 2K). During the baseline the mean degree is not different between wil types and *fmr1^-/-^* fish, but in the habituation period the *vglut2*-enriched area has higher functional connectivity than the *gad1b*-enriched area in *fmr1^-/-^* fish. This observation suggests that activity in the excitatory eurydendroid cells is more tightly coupled to brain-wide activity than the inhibitory Purkinje cells in the *fmr1^-/-^* fish.

##### Widespread hyper-excitation and loss of habenular activity in *scn1lab*

Qualitatively, *scn1lab^-/-^* animals show broader and stronger activity across the brain in response to auditory stimuli, which is less specifically located in auditory regions such as the thalamus, TS, and cerebellum compared to wild types (Figure 3A). Our voxel-based approach showed that the *scn1lab^-/-^* fish also had clear differences in the number of neurons detected in specific brain regions.

**Figure 3:**
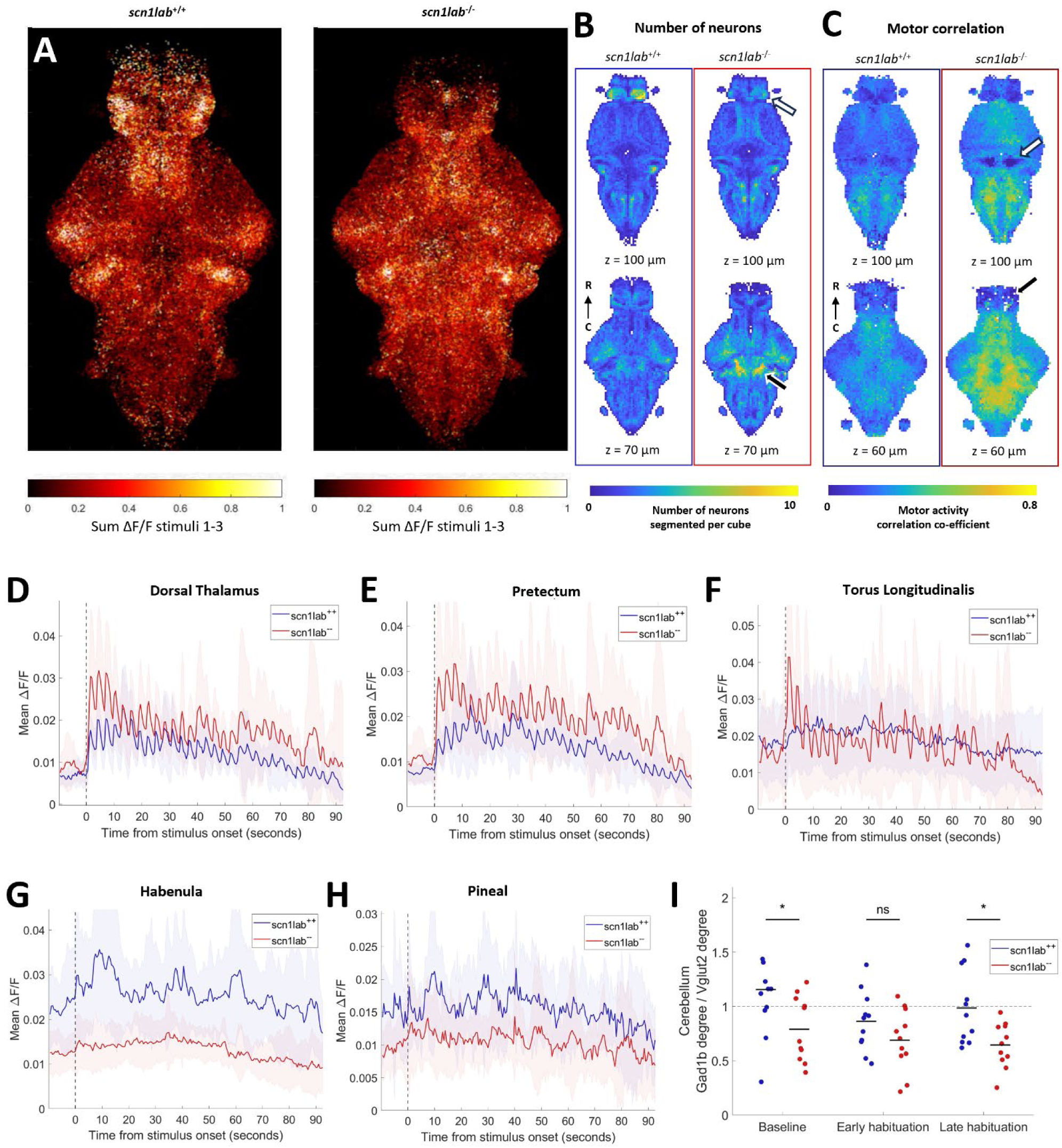
Auditory habituation phenotype in *scn1lab.* A) All segmented neurons from all fish, colored by sum of activity between stimuli 1 and 3. B) Comparison within 10 µm cubes of number of neurons segmented. Two different z-depths for each measure are shown for *scn1lab^+/+^* (left, n = 11) and *scn1lab^-/-^* (right, n = 11). Fewer neurons were segmented in the habenula (white arrow), and more in the cerebellum (black arrow) in *scn1lab^-/-^* fish than wild types. C) Comparison within 10 µm cubes of motor correlation values. Motor correlation is higher in most of the brain in *scn1lab^-/-^ fish*, except for the granule cells of the cerebellum (white arrow) and the telencephalon (black arrow). D-H) Mean activity of all neurons in the dorsal thalamus (D), pretectum (E), torus longitudinalis (F), habenula (G) and pineal (H). D-H: Shading indicates SD. I) Ratio between the degree of all neurons in the *gad1b* region of the cerebellum and the *vglut2* region of the cerebellum at different periods during habituation. Degree is based on the top 10% of edges design. Each dot represents one fish, black lines indicate the means. Significant effect of genotype (p = 0.0078), non-significant effect of time (p = 0.0567) and interaction between genotype and time (p = 0.4682, repeated measures ANOVA). The ratio is significantly lower in *scn1lab^-/-^* fish at baseline (p = 0.0437) and in the late habituation period (p = While the overall number of neurons detected brain-wide was not different (Supplementary Figure 1E), more neurons are detected throughout the mesencephalon (p = 0.0488) and rhombencephalon (p = 0.0031, including the cerebellum p = 0.0255), but fewer in the telencephalon (p = 0.0417), and notably the habenula (p = 0.0006, Figure 3B). These observations are consistent with increased proliferative cells and reduced forebrain volume of *scn1lab^-/-^* fish^21^. We also found strikingly increased motor correlations throughout most brain regions in the mutants (Figure 3C) and increased auditory correlation during the habituation period in most of these regions (Supplementary Figure 2B). The granule cells of the cerebellum deviate from this general trend, having lower correlation to both motor activity (p < 0.0001, Figure 3C) and auditory stimuli (p = 0.0006).

Analyses of cellular-resolution data in particular brain regions also reveal profound differences in the *scn1lab^-/-^* brain. Stimulus-evoked activity within the diencephalon diverges drastically from wild types, in opposing directions in different sub-regions. The *scn1lab^-/-^* fish show increased overall activity in the thalamus, particularly the dorsal thalamus, and also in the pretectum (Figure 3D and E, Supplementary Figure 2B). The y-intercept (p = 0.0219 ) and auditory correlation (p < 0.0001) are increased in the primarily glutamatergic^66^ torus longitudinalis, with clear auditory responses in *scn1lab^-/-^* but not wild types (Figure 3F). The habenula (Figure 3G) and pineal (Figure 3H), on the other hand, have drastically reduced activity in *scn1lab^-/-^* fish. In the case of the habenula this represents an almost total loss of activity. Activity is also generally decreased in the telencephalon, especially in late habituation (p = 0.0010), and decreased at baseline in the vagal ganglia (p = 0.0075) and vagal motor neuron cluster (p = 0.0205, Supplementary Figure 2B).

Similarly to *fmr1*, in *scn1lab^-/-^* fish there is diverging functional connectivity of neurons in the *gad1b* and *vglut2*-enriched areas of the cerebellum (Figure 3I), indicating imbalance between the role of excitatory and inhibitory populations.

<u>Reduced dopamine activity and changes in functional connectivity in *mecp2*</u> Unsurprisingly given its mild behavioral phenotype, *mecp2* shows only subtle changes in auditory processing and motor correlations. Indeed, a qualitative mapping of activity strength across the brain early in the stimulus train (when the behavioral phenotype is strongest) shows similar patterns across *mecp2^-/-^* larvae and their wild-type siblings (Figure 4A). There are no widespread increases in the plateau or late habituation period ΔF/F in the brain activity of *mecp2^-/-^* fish that could explain the behavioral phenotype, but there are differences in other more specific measurements that could contribute to behavior, including increases in the plateau of activity in the statoacoustic ganglion and the lobus caudalis cerebelli (Supplementary Figure 2C).

**Figure 4:**
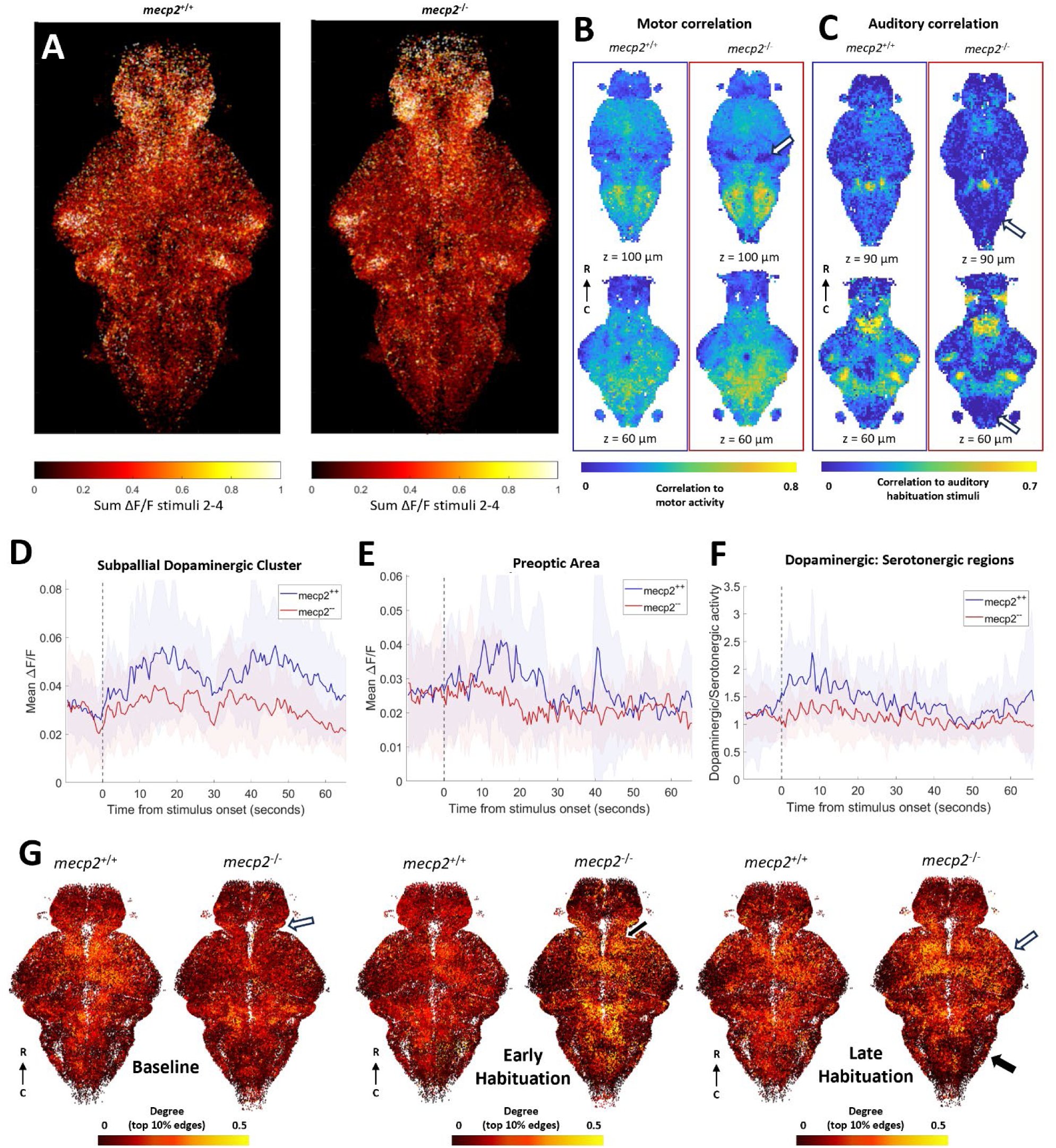
Auditory habituation phenotype in mecp2. A) All segmented neurons from all fish, colored by the sum of fluorescent activity between stimuli 2 and 4. B) Comparison within 10 µm cubes of motor correlation values. Two different z-depths for each measure are shown for *mecp2^+/+^* (left, n = 10) and *mecp2^-/-^* (right, n = 13). Motor correlations are not different in most of the brain in *mecp2^-/-^* fish, except for the granule cells of the cerebellum (white arrow). C) Auditory correlation during the habituation period is decreased in *mecp2^-/-^* fish in the inferior olive and the posterior hindbrain (white arrows). Mean activity of all neurons in the subpallial dopaminergic cluster (D)and the preoptic area (E). F) Mean ratio between activity in dopaminergic regions and serotonergic regions. D-F: Shading indicates SD. G) All neurons from all fish in a region of interest in the z-dimension. Each neuron colored by its degree as defined by the top 10% of edges during the baseline, early habituation, and late habituation periods. Differences in degree between *mecp2^-/-^* and wild-type fish are indicated in the diencephalon at baseline (white arrow), the dorsal thalamus (black arrow) during early habituation, and in the rhombencephalon (black arrow) and the mesencephalon (white arrow) during late habituation.

Looking at the motor correlations across voxels, we observe a decrease in the region of the cerebellar granule cells (p = 0.0112, Figure 4B), similar to what we observed for *scn1lab*. The correlation to auditory stimuli in the habituation period is lower in parts of the rhombencephalon, the inferior olive (p = 0.0282) and the preoptic area (p =0.0481, Figure 4C). The only difference in the sum of activity at baseline is an increase in the inferior olive in *mecp2^-/-^* fish compared to wild types (p = 0.0199, Supplementary Figure 2C).

The *mecp2^-/-^* fish have decreased activity during late habituation for neurons in dopaminergic regions such as the subpallial dopaminergic cluster (p < 0.0001 Figure 4D) and the preoptic area (p = 0.0309, Figure 4E), and a decreased number of ROIs detected in the ventral thalamus (p = 0.0169, Supplementary Figure 2C). To address the opposing effects of dopamine and serotonin on habituation, we calculated the ratio of mean activity in dopaminergic regions (preoptic area, subpallial dopaminergic cluster, pretectal dopaminergic cluster and dopaminergic cluster of the ventral thalamus) with the mean activity in serotonergic regions (superior raphe, inferior raphe and pineal).

While there is more dopaminergic than serotonergic activity in wild types, particularly at the beginning of the auditory habituation block, the mean ratio is close to 1 throughout the period for *mecp2^-/-^* fish, and this difference between genotypes is most pronounced during critical habituation period (Figure 4F).

With graph theory using the top 10% method, we observe differences in functional connectivity at the gross level of brain regions (Figure 4G). Degree is lower in *mecp2^-/-^* fish compared to wild types in the baseline period in the diencephalon (p =0.0380), and elevated in the dorsal thalamus in *mecp2^-/-^* fish in the early habituation period (p = 0.0221, Supplementary Figure 2C). In the late habituation period, high degree neurons are reduced in the rhombencephalon (p = 0.0156) and increased in the mesencephalon (p =0.0326) in the *mecp2^-/-^* fish compared to wild types.

##### Increased activity and disrupted functional connectivity in *cntnap2*

The *cntnap2a^-/-^b^-/-^* double knockouts have an (insignificantly) elevated y-intercept, slower habituation, and a higher plateau compared to wild-type siblings (Figure 2G). When these combined effects are strongest, during early habituation, neurons across the brain are more responsive to auditory stimuli (Figure 5A). The *cntnap2a^-/-^b^-/-^* larvae have several brain regions with increased y-intercepts, including most of the hindbrain, but fewer that have differences in decay rate or plateau (Supplementary Figure 2D, Figure 5B). There is also increased correlation to auditory stimuli during the habituation period in the preoptic area, subpallium, and eminentia thalami (thalamic eminence) (Figure 5C).

**Figure 5:**
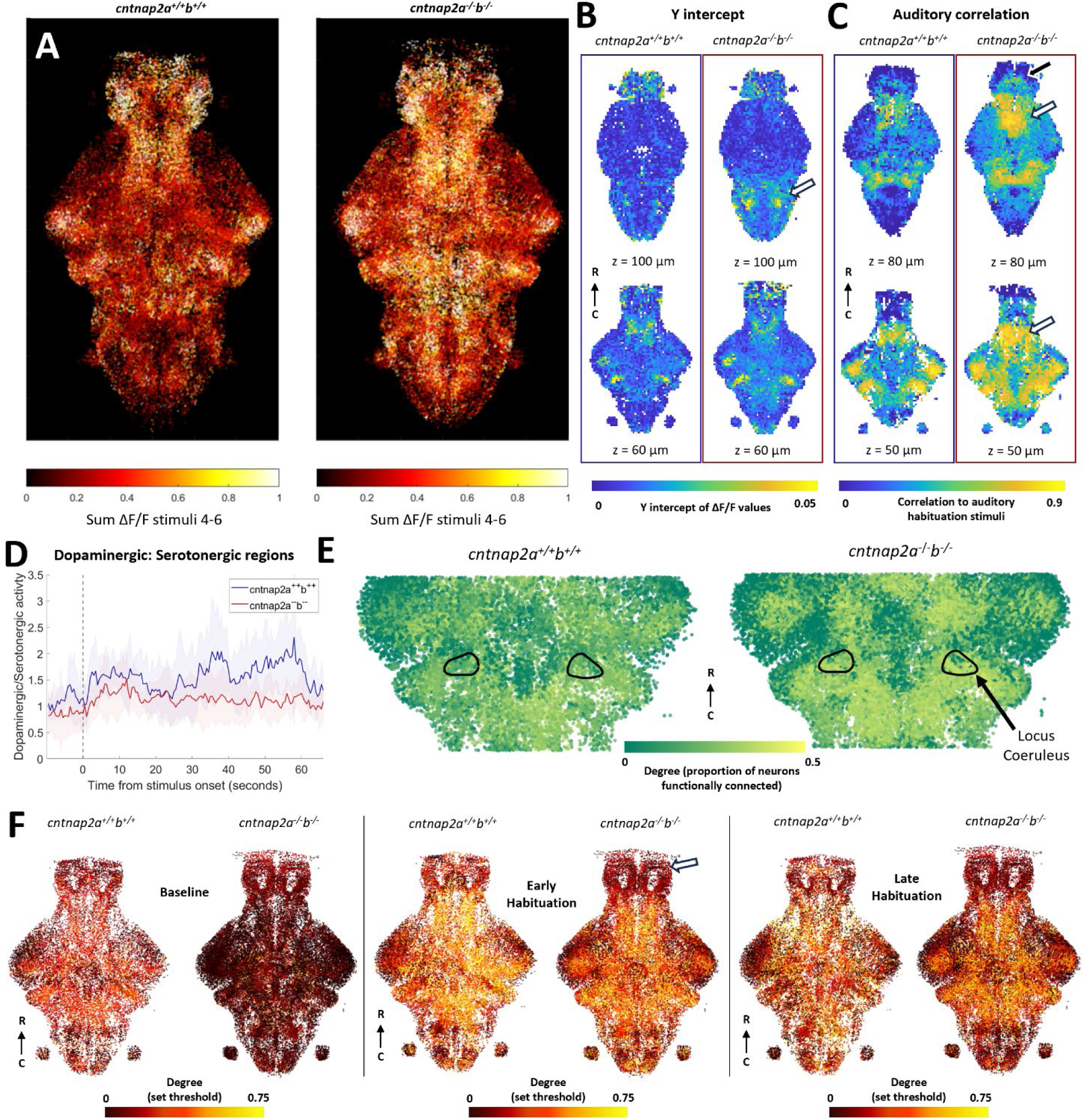
Auditory habituation phenotype in *cntnap2*. A) All segmented neurons from all fish, colored by the sum of fluorescent activity between stimuli 4 and 6. B) Voxel-based representations of curve fits during habituation period. Two different z-depths for each measure are shown for *cntnap2a^+/+^b^+/+^* (left, n = 5) and *cntnap2a^-/-^b^-/-^* (right, n = 7). Y-intercepts are higher in *cntnap2a^-/-^b^-/-^* fish than wild types in the pineal and most of the rhombencephalon (white arrow). C) Voxel-based auditory correlation values during the habituation period. Auditory correlation is not different in most of the brain in *cntnap2a^-/-^b^-/-^* fish, except for the subpallium (black arrow), preoptic area (white arrows), and pineal. D) Mean ratio of activity in dopaminergic regions versus serotonergic regions. Shading indicates SD. E) Sub-selection of neurons, colored by degree, as determined using the top 10% of edges from correlation during the whole habituation period. Black outlines indicate the locus coeruleus. F) All neurons from all fish in a section of interest between 50-75 µm depth in the z-dimension. Each neuron is colored by its degree as defined by a set correlation threshold during the baseline, early habituation, and late habituation periods. At baseline, neurons have lower degree throughout the brain in the *cntnap2a^-/-^b^-/-^* fish compared to wild types. In the early habituation period, only the telencephalon has lower degree *cntnap2a^-/-^b^-/-^* fish compared to wild types (white arrow).

We observed both decreased activity in dopaminergic regions (preoptic area, thalamic dopaminergic cluster, tegmentum and subpallial dopaminergic cluster) and increased activity in serotonergic regions (superior raphe and pineal, Supplementary Figure 2D). While the ratio between dopaminergic activity and serotonergic activity in the mutants (as calculated for *mecp2*) shows a similar increase at the beginning of the stimulus train to that of the wild types, it remains below the mean value for wild types until the end of the auditory stimuli (Figure 5D). This suggests that the temporal dynamics of this ratio are preserved at the onset of the sound, but not in the later part of the train, when the behavioral phenotype is strongest.

According to graph theory measurements with the top 10% thresholding method, functional connectivity in early habituation in *cntnap2a^-/-^b^-/-^* fish is decreased across several parts of the forebrain, and consequently increased in the rhombencephalon (p = 0.0197). The locus coeruleus also has higher functional connectivity during the whole period (p = 0.0001, Figure 5E). With the hard threshold method, functional connectivity is broadly reduced at baseline (Figure 5F). This effect is not due to differences in motor activity at baseline: both motor activity (p = 0.0029) and genotype (p = 3.08 x10^-6^) have significant effects on the threshold required to attain the top 10% of correlations (linear mixed-effect model). The decreased functional connectivity persists in the telencephalon (p = 0.0008) and parts of diencephalon into early but not late habituation (Figure 5F). The few regions without lower degree during the baseline include monoaminergic regions such as the subpallium, preoptic area, locus coeruleus, and superior raphe (Supplementary Figure 2D). Fish mutant for only *cntnap2a* or *cntnap2b* generally resemble the double mutants, but the mutant phenotype diverges between the single mutants in the functional connectivity of the locus coeruleus and the set threshold functional connectivity over time (Supplementary Figure 4).

## Discussion

### Unique phenotypic fingerprints for each gene

In this study, we have used auditory habituation as a paradigm to characterize the behavioral and brain-wide phenotypes for four genetic lines with relevance to autism. Each gene showed a different combination of traits describing its behavioral phenotype and each had its own profile of activity changes across the brain. The points of overlap, but also the distinctions between the lines’ phenotypes, raise interesting questions about the various ways in which changes in brain activity could lead to altered perception and behavior. This approach is a first step toward understanding the diverse and multigenic ways in which sensation and behavior are altered across the autism spectrum in humans.

#### Initial hyperresponsiveness in *fmr1*

The behavioral phenotype of *fmr1* was limited to an increase in the initial responsiveness (Figure 1D). Correspondingly, we found increased initial response amplitude in several auditory brain structures, including the SAG, torus semicircularis, and thalamus, and auditory neurons in the ON. We therefore postulate that the auditory hypersensitivity phenotype arises within the auditory pathway, which drives Surprisingly, we also found some differences in the late habituation period, despite the lack of a behavioral phenotype in the plateau of responses. The plateau of several rhombomeres of the hindbrain is also decreased in *fmr1^-/-^* fish, suggesting lower activity in motor output regions. Indeed, there is also increased activity in the granule cells of the cerebellum in the early part of the habituation period (Figure 2), which may represent more inhibition of motor output^67–70^. The *fmr1^-/-^* larvae may therefore undergo stronger adaptation to reach the same behavioral plateau as the wild types, having started from a greater initial response.

While the hypersensitivity fits with previous zebrafish *fmr1* studies^32^, the lack of a phenotype in the habituation to sounds here is at odds with what would be expected from studies of auditory habituation in FXS^8^. It is also different to the decreased auditory adaptation in the brain activity of *Fmr1^-/-^* mice, although different age or stimulus presentation rates may explain this discrepancy^71^.

#### Drastically increased responsiveness in *scn1lab*

The behavioral phenotype in *scn1lab* was the strongest of the four genes, with highly elevated initial response and plateau responses (Figure 1E). Unsurprisingly, we also found the most dramatic differences in brain activity in *scn1lab* mutants (Supplementary Figure 2B). Because there are differences in the volume and number of proliferative cells in the brains of *scn1lab^-/-^* fish^21^, we infer that neuronal proliferation, migration, differentiation and/or survival is altered in the brain. These changes likely lead to some neurons playing different roles within the network, such as those in the torus longitudinalis responding completely differently in *scn1lab^-/-^* animals than wild types (Figure 3F).

We observed both increases and reductions in activity in distinct parts of the brains of *scn1lab^-/-^* fish. Activity was increased in the thalamus, pretectum, tegmentum, and torus longitudinalis. Conversely, activity was reduced in the telencephalon, habenula, pineal, and vagal ganglia. A previous study using pERK/tERK staining to infer activity levels found decreased activity at baseline throughout the brains of *scn1ab^-/-^* fish, though most strikingly in the telencephalon and habenula^21^. Our results recapitulate this forebrain phenotype, but also uncover various changes in activity across the rest of the brain.

The habenula has a role in suppressing anxiety or fear responses in zebrafish ^43,72–75^. The putative ‘induced passivity’ network involves increased activity in the habenula and decreased activity in the dorsal thalamus ^43^. The loss of activity in the habenula and increased activity in the dorsal thalamus we observe in *scn1lab^-/-^* fish may therefore represent an ‘anti-passivity’ network, which leads to greatly increased behavioral output. However, our methods only measure activity of neurons, and glia have also been shown to be important for passivity^59^.

The higher correlation to motion throughout the brains of *scn1lab^-/-^* animals (Figure 3C) may be due to greater physical movement in *scn1lab^-/-^* fish, which recruit brain-wide activity more than the comparatively smaller movements of the wild types. An alternate explanation is functional hyperconnectivity across the mutant brain, which would tie to the role for *scn1lab* in epilepsy^76^. However, we do not observe gross differences in graph theory measures of network connectivity.

#### Higher plateau of responses in *mecp2*

The behavioral phenotype for *mecp2* was specifically an increase in the plateau of the response rates, suggesting decreased habituation (Figure 1F). A recent study measured auditory habituation in *mecp2^- /-^* zebrafish larvae, and found no differences in the habituation as measured by the likelihood of startle^77^. It is not clear whether this study may have found an increase in the plateau of response rates using the distance travelled, as we have here, instead of startle probability. In our study, there were no differences in pure activity level in auditory regions to explain this phenotype. The only regions with a higher plateau of responses were the SAG and lobus caudalis cerebelli, which is generally regarded to be part of the vestibular network^61^. The vestibular and auditory systems are functionally intertwined in larval zebrafish, however, and these higher plateaus may therefore correspond to altered auditory, rather than vestibular processing^78^.

We also observe differences in the functional connectivity of the brain networks over time in the major divisions of the brain (Figure 4G). In the early part of habituation period, where there are no behavioral phenotypes, only the dorsal thalamus has higher functional connectivity in *mecp2^-/-^* fish. By the end of the habituation period, when the behavioral phenotype emerges, the balance of edges in the network is shifted toward the mesencephalon and away from the rhombencephalon in *mecp2^-/-^* fish. This matched timing with the behavioral phenotype suggests it may have functional consequences for the behavior of the animal. In contrast to *fmr1* and *scn1lab*, where an initial hypersensitivity either is (*fmr1*) or is not (*scn1lab*) compensated for during habituation, *mecp2* provides an example of a genetic line with a normal initial response that only exhibits a phenotype as habituation plays out. It will be interesting to explore whether the earlier divergence of functional connectivity in the dorsal thalamus may be linked to the ensuing shift in functional connectivity across the brain.

#### Slower habituation and higher plateau in *cntnap2*

In the behavior of *cntnap2a^-/-^b^-/-^* fish, we observed a slower decay rate and a higher plateau of responses to auditory stimuli (Figure 1G). Surprisingly, given that there was no significant behavioral difference in the initial response, we found far more differences in the y-intercept of brain activity than either the decay rate or plateau (Supplementary Figure 2D). These increases occurred throughout the rhombencephalon, as well as in the pineal and subpallial dopaminergic cluster. Our best explanation for this discrepancy is that, given the difficulty of getting a large experimental n for a duplicated gene such as *cntnap2*, we lacked statistical power to identify what is a real y-intercept phenotype, yielding only an insignificant trend (Figure 1G).

We also observed widespread decreased functional connectivity throughout the brain in *cntnap2a^-/-^b^-/-^* fish at baseline, independent of differences in motor activity (Figure 5F). In the early habituation period, these differences disappear except in the telencephalon and habenula, and by the late habituation period, the network connectivity is comparable to that in wild types. This initial lack of functional connectivity in the network, and late recruitment of higher-order integrative structures, may underlie the reduced ability of the network to adapt to auditory stimuli over time. Interestingly, each of the single mutants resembled one aspect of these functional connectivity phenotypes (Supplementary Figure 4H). Each mutation may therefore contribute differently to the cumulative phenotype of the double mutant.

Similarly to *scn1lab^-/-^* regions outside of the forebrain, we did not observe widespread changes in activity at baseline as reported in a previous paper that used pERK/tERK staining to infer brain activity^21^ Our baseline is measured over a fairly short period preceding stimuli, whereas the increased pERK/tERK activity may be related to freely swimming in the environment over a longer period. The observed decrease in habituation to auditory stimuli does, however, fit with previous studies in *Cntnap2* knock-out rats^79,80^.

### Overlapping phenotypes across genetic lines

There were not overlapping phenotypes between genes in the primary auditory regions, rather shared circuitry changes appear to be in the sensorimotor and modulatory regions.

The y-intercept of neurons in rhombomeres 3-5 is higher in *scn1lab^-/-^* and *cntnap2a^-/-^b^-/-^* fish than wild types, which likely relates to activity in motor regions driving the increased initial behavioral response in each of those lines. Conversely, the plateau of activity in this part of the hindbrain is decreased in *fmr1^-/-^* fish, which supports the idea that the network is more strongly adapted from a higher initial point to reach the same plateau as wild types.

The reduced correlation to motor activity in the granule cells of the cerebellum is a striking phenotype in *scn1lab^-/-^* and *mecp2^-/-^* fish (Figure 6), particularly because it is scaled to the intensity of the behavioral phenotype of each gene. Previous studies in larval zebrafish suggest suppression of granule cell activity is required for stimulus-evoked behavioral responses and preventing immobility ^67–70^.

**Figure 6:**
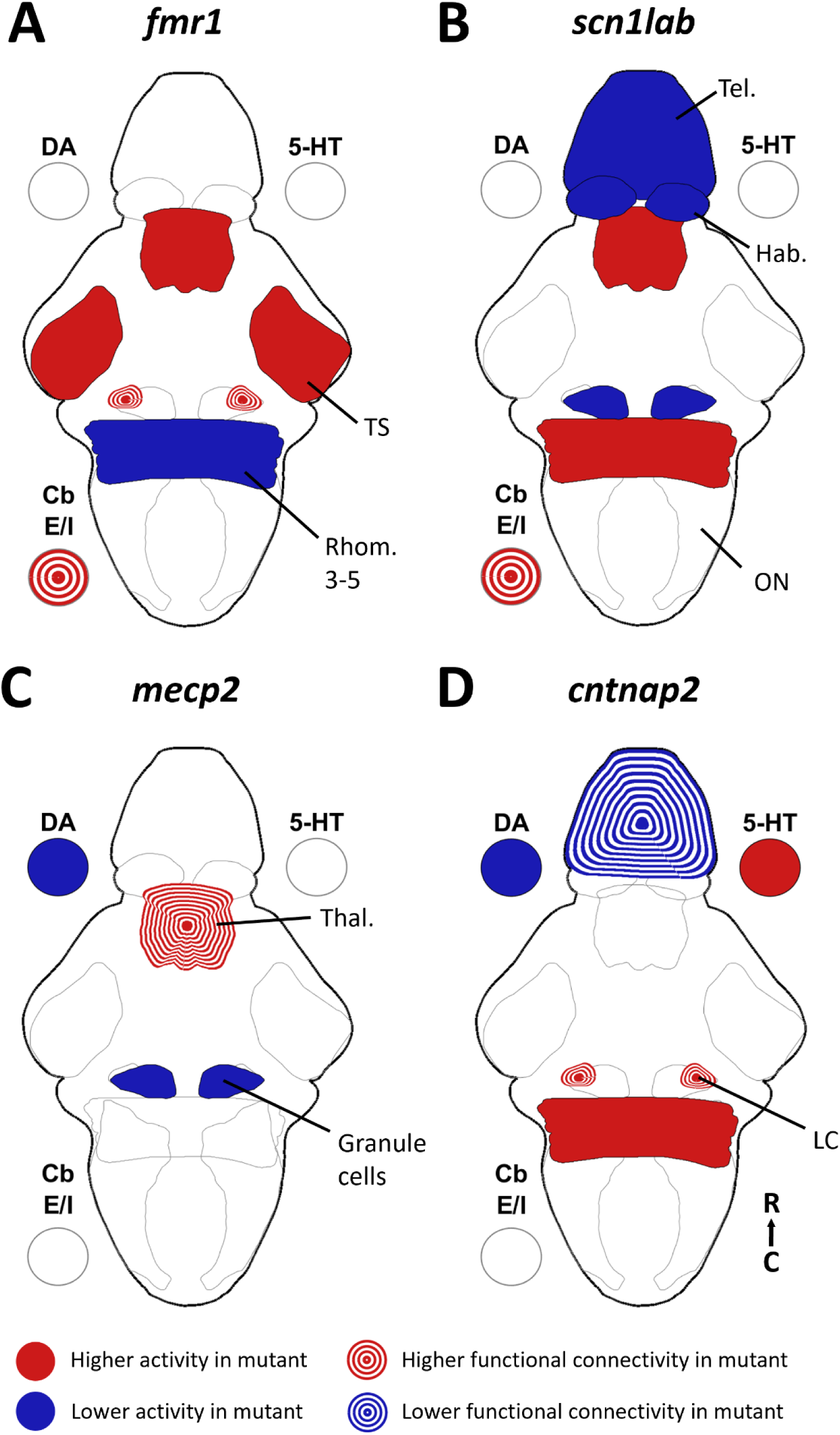
Summary of brain activity underlying auditory habituation phenotypes. A summary of the brain-wide phenotypes for activity and functional connectivity found in *fmr1* (A), *scn1lab* (B), *mecp2* (C), and *cntnap2* (D). 5-HT = serotonin, DA =dopamine, Cb E/I = cerebellum excitatory/inhibitory regions, Hab = habenula, LC = locus coeruleus, ON = octavolateralis nucleus, Rhomb. = rhombomere, Tel. = telencephalon, Thal = thalamus, TS = torus semicircularis.

Reduced granule cell activity in *scn1lab* and *mecp2* mutants at the time of stimulus-evoked movements may therefore contribute to sustained higher behavioral responses in both these lines.

We found that the functional connectivity of the *gad1b*-enriched and *vglut2*-enriched parts of the cerebellum (as measured by the degree) diverged similarly in both *fmr1* and *scn1lab* mutants (Supplementary Figure 2,Figure 6). While there is some heterogeneity of cell types within these regions, we take these populations to be representative of the Purkinje cells (*gad1b*) and eurydendroid cells (*vglut2*). The functional connectivity of the eurydendroid cells is higher than that of the Purkinje cells in both the *fmr1* and *scn1lab* mutants, particularly in the late habituation period. This divergence in connectivity of excitatory and inhibitory regions may relate to the theory of E/I imbalance in autism^12^. Considering the crucial role of the cerebellum in sensorimotor processing, this divergence in connectivity seems likely to affect behavioral output. It is particularly interesting for *fmr1* that this connectivity phenotype is strongest in the late habituation period when there is not a behavioral phenotype in the plateau of responses, indicating that while the behavioral plateau reached resembles that of the wild types, the underlying brain activity is not the same.

The *cntnap2a^-/-^b^-/-^* and *mecp2^-/-^* fish both have differences in monoaminergic activity (Figure 6), which fit with what we may expect based on pharmacological studies ^38,39^. Indeed, pharmacological manipulations of dopamine and serotonin receptors affected baseline activity in a different *cntnap2a^-/-^ b^-/-^* line^48^. The combination of dopaminergic and serotonergic phenotypes, and the earlier onset of these differences in *cntnap2,* may explain why the behavioral output of these fish diverges from wild types earlier in the stimulus train than in *mecp2* fish, which have only dopaminergic differences. The difference in ratio of dopaminergic to serotonergic activity was recapitulated in both of the single gene *cntnap2* mutants (Supplementary Figure 4E). There are also some reductions in activity in dopaminergic areas of forebrain in *scn1lab^-/-^* fish, but these are less likely to be dopamine-specific given the reduced number of dopaminergic forebrain neurons in *scn1lab^-/-^* fish^21^ and the overall reduction in telencephalic activity (Supplementary Figure 2). Interestingly, the dopaminergic effects are specific to the subpallium and preoptic area, and the serotonergic differences in *cntnap2* are stronger in the pineal than the superior raphe, the serotonergic region typically associated with habituation^38^.

Functional connectivity over the full habituation period is increased in the locus coeruleus in both *fmr1* and *cntnap2* fish (Figure 6). The locus coeruleus is a noradrenergic center, whose activity is associated with alertness^81^. This increased recruitment of the locus coeruleus within the brain network may indicate a generally more alert state in these fish, leading to hyperresponsiveness to auditory stimuli. We also observe an increased plateau of activity in the locus coeruleus in *scn1lab^-/-^* fish (Supplementary Figure 2), which may represent more persistent alertness late in the habituation period, consistent with the observed behavior.

We observe differences in the activity or functional connectivity of parts of or the whole telencephalon in all three of the genes that have higher plateau of responses: *mecp2*, *scn1lab* and *cntnap2* (Figure 6). The telencephalon is the seat of higher-order processing in the zebrafish brain^82,83^, so it is unsurprising that it would have involvement in adaptation to stimuli. It is notable that *fmr1*, the only gene not to have a behavioral phenotype in the plateau of responses, does not have significant changes to the activity of the telencephalon.

### Future directions

We have identified a range of differences in brain activity in these four autism-relevant genetic lines. In each, we have addressed brain-wide function in a fundamental way, observing activity across the entire brain at cellular resolution. Doing so has permitted several separate and complementary approaches for assessing brain-wide activity and the ways in which networks may change differently in different autism-relevant mutant lines.

This approach, in its current form, comes with important limitations, especially because we cannot assess the synaptic relationships between the neurons that we observe and cannot conclusively determine the neurons’ neurotransmitter subtypes. As such, while these results are comprehensive from one perspective, they represent merely a departure point for better understanding the mechanisms by which information flow changes in the brains of these fish. Future studies could shed greater light on these changes by performing calcium imaging while co-labelling neurons with transgenic markers for neurotransmitter subtypes, such as the GABAergic, glutamatergic, dopaminergic, and serotonergic populations that our study implicate. Further, optogenetic techniques could be employed to manipulate parts of the circuit in wild types that we have implicated in our mutants, directly testing the hypothesized impacts on activity elsewhere in the brain and on behavior. More in-depth behavioral analysis may also provide more insight into brain activity underlying changing response strategies.

Additionally, it would be interesting to add more genes to this collection. Here we have presented four different autism-associated genes, but this only scratches the surface of autism’s genetic complexity and phenotypic diversity. Adding more lines with different combinations of behavioral and neural phenotypes would enable a greater understanding of which combinations of brain activity phenotypes link to which behaviors. Ultimately, this will lead to a fuller appreciation of the relationship among genetics, perception, and behavior.

## Supporting information

Supplementary Information: all p-values for grid comparisons in Supplementary Figure 2

## Author Contributions

Experimental design: MW, AG, REP, LC, EJH and EKS. Experimental methodology: MW, REP, IAF, TJK, LAS, and CL. Data collection: MW, AG, TM, AM, and LC. Processing of calcium imaging data: MW, JA and WQ. Processing of behavioral data: LAS, CL and TM. Data analysis: MW and CL. Funding secured by: EKS, IAF, CL, EJH and SP. Writing: MW and EKS. Supervision: EKS. All authors read and approved the final manuscript.

## Funding

Support was provided by a Simons Foundation Research Award (625793), two ARC Discovery Project Grants (DP220103812 and DP230102614), and an NHMRC Investigator Grant (2027072) to EKS. The research reported in this publication was supported by the National Institute of Neurological Disorders and Stroke of the National Institutes of Health under Award Number R01NS118406 to EKS. The content is solely the responsibility of the authors and does not necessarily represent the official views of the National Institutes of Health. Support was also provided by an ARC DECRA award (DE220100691 & DE230100972) to CL and IAF, an NHMRC Ideas Grant (2012140) to IAF and a Simons Foundation Research Award (573508) to EJH. MW is supported by a University of Queensland RTP scholarship.

## Acknowledgements

The authors would like to acknowledge the University of Queensland’s Biological Resources aquatics team and the *Danio rerio* University of Melbourne facility (DrUM, Melbourne, Australia) for maintenance of zebrafish lines, the Queensland Brain Institute workshop staff for 3D-printing of imaging chambers, and Dr Summer Thyme for her feedback on the manuscript.

## Ethics approval

All work was performed in accordance with research application SBS/341/19 and breeding application IMB/271/19/BREED, which was approved by the Anatomical Biosciences Animal Ethics Committee at the University of Queensland, or research application 2022-24987-35220-5, which was approved by the SLA-2 Animal Ethics Committee at the University of Melbourne.

## Notes

### Competing Interest Statement

The authors have declared no competing interest.

