## Supplementary Information: Supplementary Table 1 and Supplementary Figures 1-4 for "Brain-wide circuitry underlying altered auditory habituation in zebrafish models of autism"

*Supplementary table 1: Primer sequences used for PCR genotyping of experimental animals in autism auditory habituation experiments.*

| **Gene** | **Forward primer sequence** | **Reverse primer sequence** |
|---|---|---|
| *fmr1* | 5' – CTA AAT GAA ATC GTC ACA TTA GAG | 5' – TCC ATG ACA TCC TGC ATT AG |
| *scn1lab* | 5' – CAG CAA ATA AAT GAA CGC CTT A | 5' – AGA GAG TTA CCA CAA ACA CAC TCG |
| *mecp2* | 5' – AAA GGA AAG GCA TGA TGT GG | 5' – ATA CAT TGG GCC TCT GTC CC |
| *cntnap2a* | 5' – ACC CTT AAA ATT GAT AAA AGA ACA CG | 5' – GCA GAA AAG GGG CTA AAT TAA AA |
| *cntnap2b* | 5' – TGC GAT GTG TAT CAT ATG TTC TTT T | 5' – AAA AAG GTA GCT CAA ACT GTA ATT G |

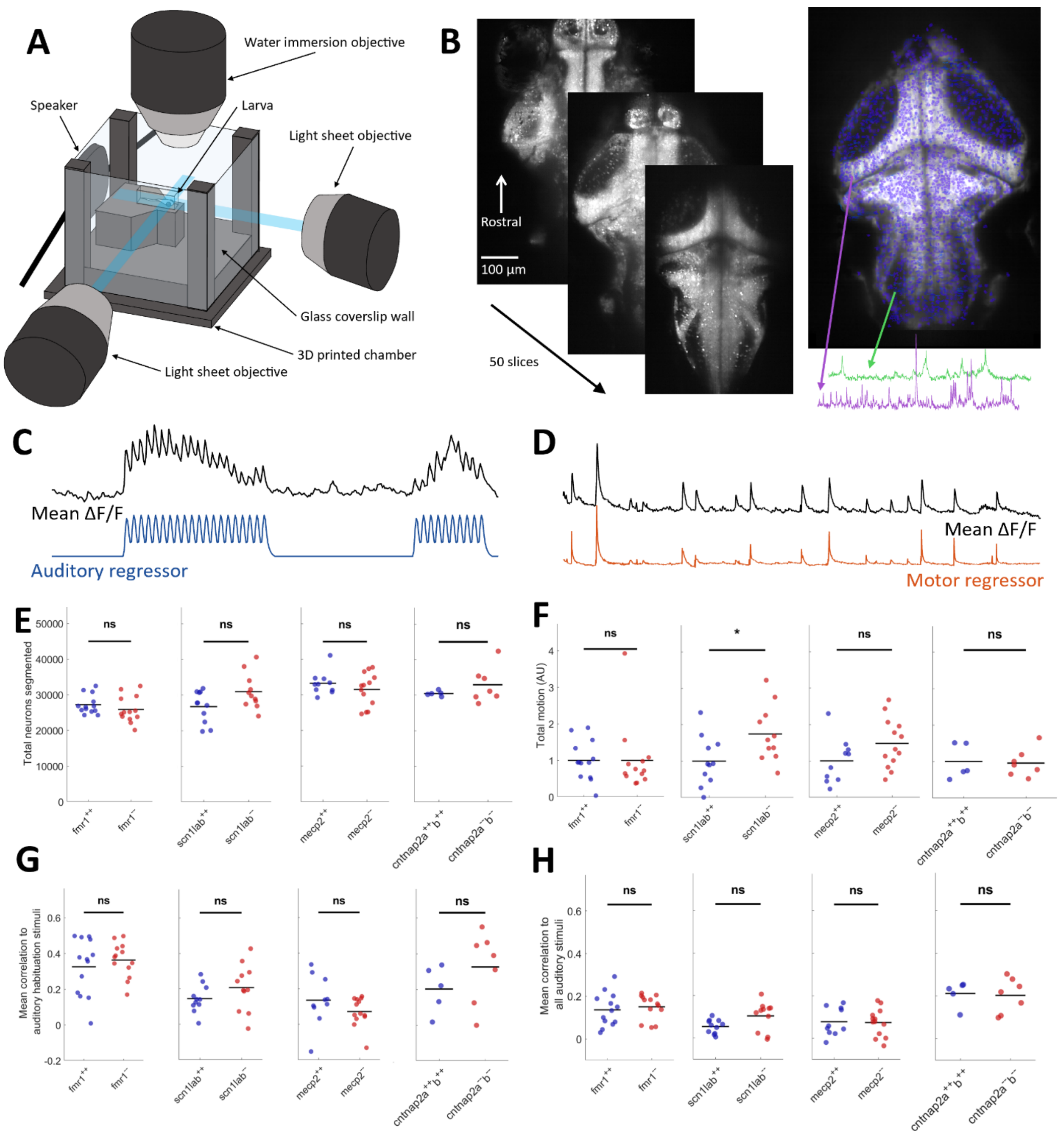


Supplementary Figure 1: Imaging of brain-wide neuronal activity during auditory habituation. A) Selective plane illumination microscope set-up. Larva mounted in low melting point agarose is illuminated by two perpendicular sheets of light. Acoustic stimuli are delivered by a speaker affixed to the back wall. Calcium activity is captured with a water-immersion objective. B) Image processing pipeline. 50 z-planes spanning the dorsoventral axis are acquired at a rate of 2Hz. Individual neurons are segmented using Suite2p, and their fluorescent traces extracted (examples shown in green and purple). C) Example auditory regressor and mean trace from auditory-classified neurons from a single fish. D) Example motor regressor and mean trace from motor-responsive neurons from a single fish. E) Total number of neurons segmented from each fish in each dataset. F) Total area under the curve of the motor regressor, normalized to the mean value for wild types of each dataset. * = $p < 0.05$. G) Mean correlation to the auditory stimuli in the habituation period from all neurons. H) Mean correlation to all auditory stimuli. For E-H, each dot represents one fish, black lines show means.

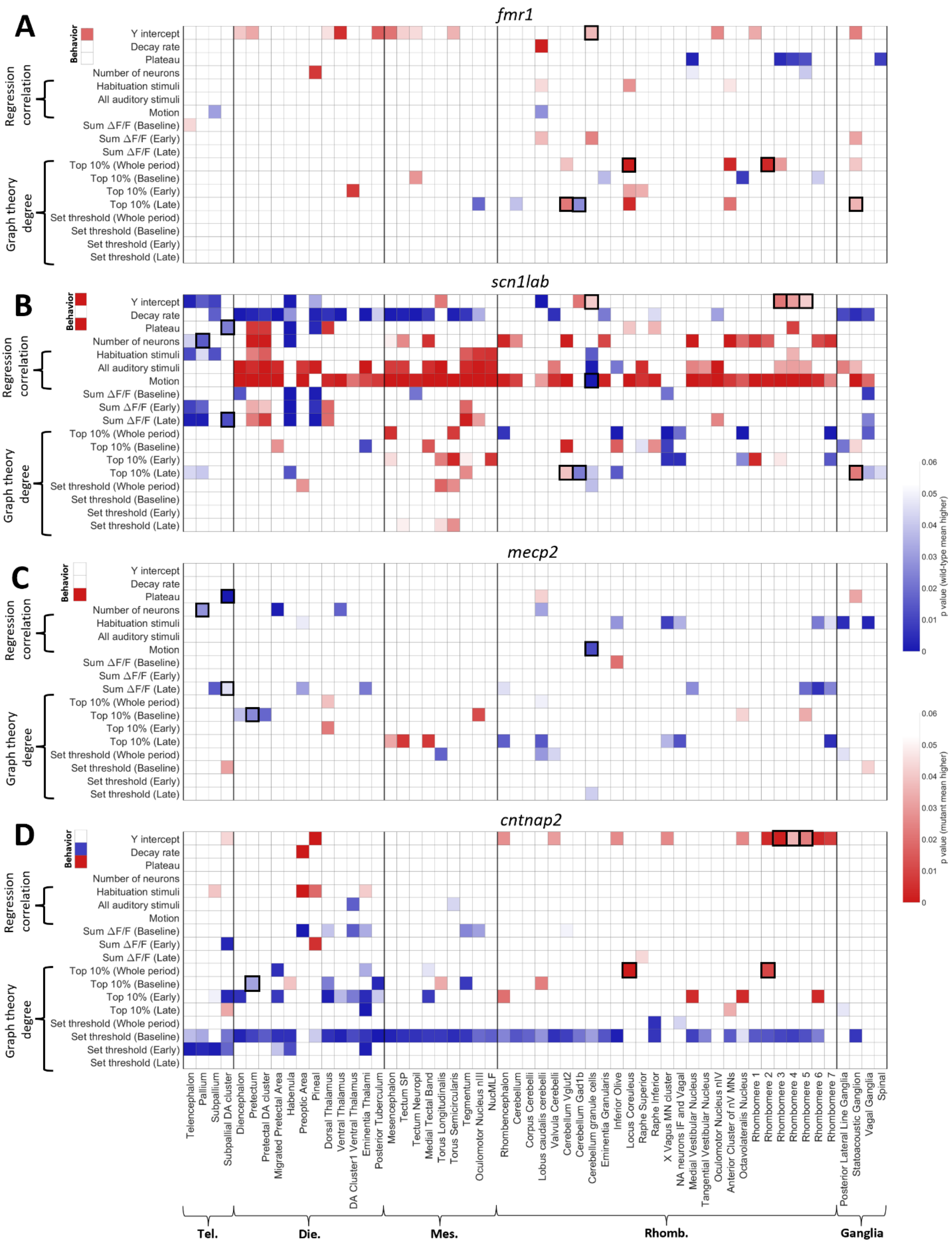
A
fmr1
B
scn1lab
C
mecp2
D
cntnap2
Behavior
Y intercept
Decay rate
Plateau
Regression correlation
Number of neurons
Habituation stimuli
All auditory stimuli
Motion
Sum ΔF/F (Baseline)
Sum ΔF/F (Early)
Sum ΔF/F (Late)
Graph theory degree
Top 10% (Whole period)
Top 10% (Baseline)
Top 10% (Early)
Top 10% (Late)
Set threshold (Whole period)
Set threshold (Baseline)
Set threshold (Early)
Set threshold (Late)
p value (wild-type mean higher)
p value (mutant mean higher)
Telencephalon
Pallium
Subpallium
Subpallial DA cluster
Diencephalon
Pretectum
Pretectal DA cluster
Migrated Pretectal Area
Habenula
Preoptic Area
Pineal
Dorsal Thalamus
Ventral Thalamus
DA Cluster1 Ventral Thalamus
Eminentia Thalami
Posterior Tuberculum
Mesencephalon
Tectum SP
Tectum Neuropil
Medial Tectal Band
Torus Longitudinalis
Torus Semicircularis
Tegmentum
Oculomotor Nucleus nIII
NucMLF
Rhombencephalon
Cerebellum
Corpus Cerebelli
Lobus caudalis cerebelli
Valvula Cerebelli
Cerebellum Vglut2
Cerebellum Gad1b
Cerebellum granule cells
Eminentia Granularis
Inferior Olive
Locus Coreuleus
Raphe Superior
Raphe Inferior
X Vagus MN cluster
NA neurons IF and Vagal
Medial Vestibular Nucleus
Tangential Vestibular Nucleus
Oculomotor Nucleus nIV
Anterior Cluster of nV MNs
Octavolateralis Nucleus
Rhombomere 1
Rhombomere 2
Rhombomere 3
Rhombomere 4
Rhombomere 5
Rhombomere 6
Rhombomere 7
Posterior Lateral Line Ganglia
Statoacoustic Ganglion
Vagal Ganglia
Spinal
Tel.
Die.
Mes.
Rhomb.
Ganglia

Supplementary Figure 2: Summary graphs of comparisons of different metrics across different brain regions using linear mixed effects models. The same comparisons are presented for each of the four genes. Grid locations are colored by the p-value of the effect of genotype, all p > 0.05 are white. Red colored squares indicate the metric is significantly higher in the mutant, blue indicates the metric is significantly higher in the wild type. Full p values can be found in supplementary material. Dark outlines indicate phenotypes present in two genes. Insets display the p-values from curve fit comparisons of behavioral data. DA = dopaminergic, NA = noradrenergic. SP = Stratum Periventriculare, Tel. = Telencephalon, Die. = Diencephalon, Mes. = Mesencephalon, Rhomb. = Rhombencephalon. MN = motorneuron. NucMLF = Nucleus of the medial longitudinal fascicle. IF = Interfascicular

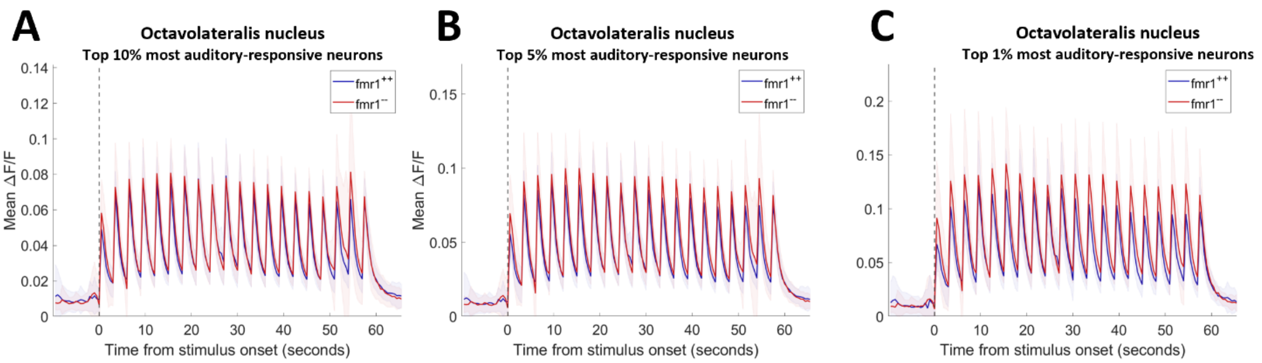


Supplementary Figure 3: Mean traces of neurons in the ON of the *fmr1* group, with increasing threshold stringency for correlation to all auditory stimuli. Mean traces are shown for the top 10% (A), 5% (B) and 1% (C) of auditory neurons in the ON of each fish. Shading indicates std.

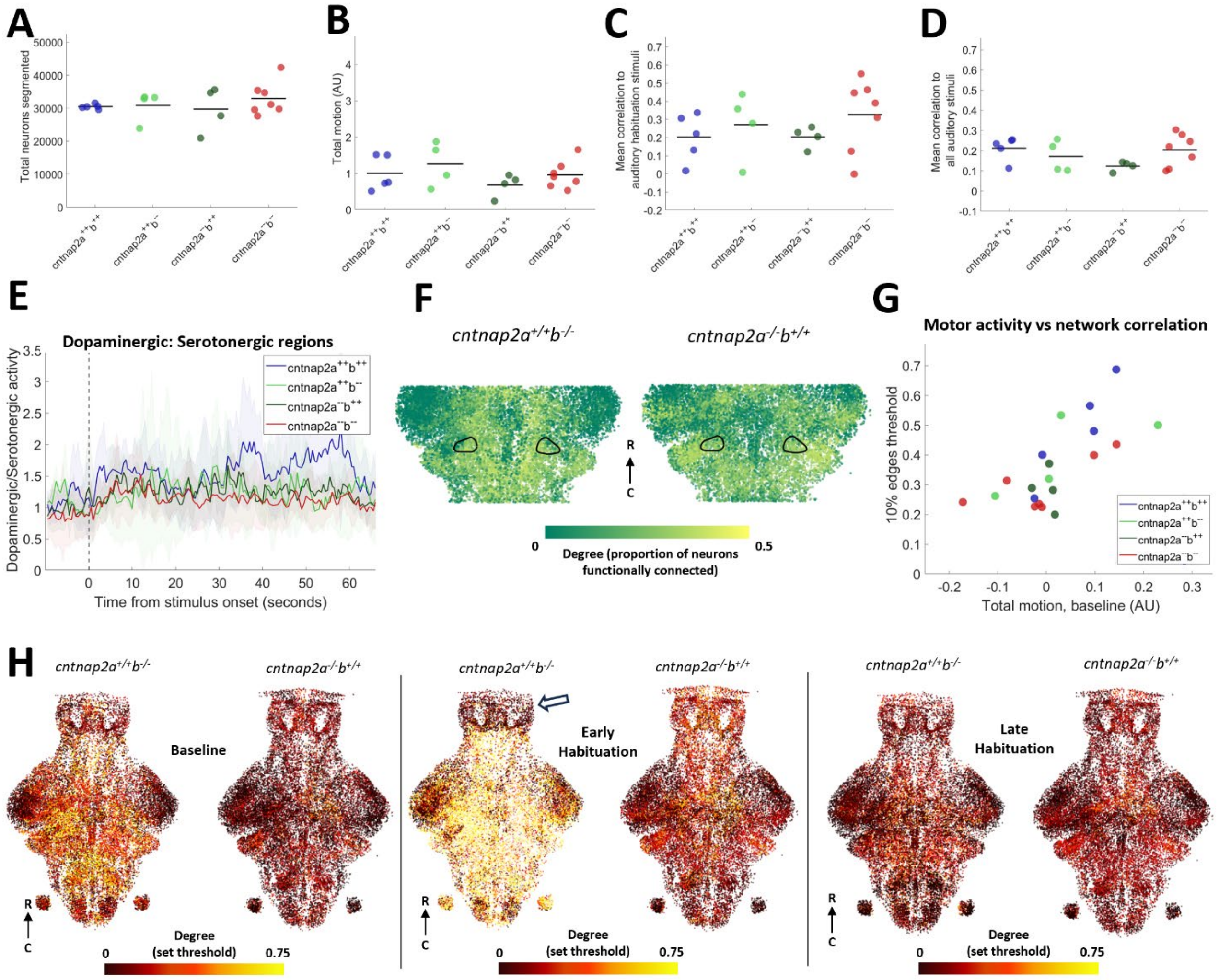

Supplementary Figure 4: Auditory habituation in *cntnap2* single mutants. A) Total number of neurons segmented from each fish in each dataset. B) Total area under the curve of the motor regressor, normalized to the mean value for wild types of each dataset. C) Mean correlation to the auditory stimuli in the habituation period from all neurons. D) Mean correlation to all auditory stimuli. For A-D, each dot represents one fish, black lines show means. E) Mean ratio between activity in all dopaminergic regions and all serotonergic regions. Shading indicates SD. F) Sub-selection of neurons, colored by their degree, as determined using the top 10% of edges from correlation during the whole habituation period. Black outlines indicate the locus coeruleus. G) Sum motor activity compared to correlation threshold required for the top 10% of edges during the baseline period. Each dot represents one fish. Linear mixed-effect model indicates the amount of motor activity affects the overall correlation throughout the brain ($p = 0.0004$), but the effect of genotype is also strong ($p < 0.0001$). H) All neurons from all fish in a region of interest in the z-dimension, between 50 – 75 µm depth. Each neuron colored by its degree as defined by a set correlation threshold during the baseline, early habituation period and late habituation period.
